# Mesenchymal Meis2 controls whisker development independently from trigeminal sensory innervation

**DOI:** 10.1101/2024.08.13.607774

**Authors:** Mehmet Mahsum Kaplan, Erika Hudacova, Miroslav Matejcek, Haneen Tuaima, Jan Krivanek, Ondrej Machon

## Abstract

Hair follicle development is initiated by reciprocal molecular interactions between the placode-forming epithelium and the underlying mesenchyme. Cell fate transformation in dermal fibroblasts generates a cell niche for placode induction by activation of signaling pathways WNT, EDA, and FGF in the epithelium. These successive paracrine epithelial signals initiate dermal condensation in the underlying mesenchyme. Although epithelial signaling from the placode to mesenchyme is better described, little is known about primary mesenchymal signals resulting in placode induction. Here we show that *Meis2* expression in cells derived from the neural crest is critical for whisker formation, and also for branching of trigeminal nerves. While whisker formation is independent of the trigeminal sensory innervation, MEIS2 in mesenchymal dermal cells orchestrates the initial steps of epithelial placode formation and subsequent dermal condensation. MEIS2 regulates the expression of transcription factor *Foxd1*, which is typical of pre-dermal condensation. However, deletion of *Foxd1* does not affect whisker development. Overall, our data suggest an early role of mesenchymal MEIS2 during whisker formation and provide evidence that whiskers can normally develop in the absence of sensory innervation or *Foxd1* expression.

## Introduction

The local cell environment provides essential cues that determine cell fate decisions, proliferation, and migration. The molecular communication between the developing epithelium and the underlying cell environment is crucial for the formation of multiple structures, which are derived from the embryonic epithelium, such as the eye lens, hair follicles including whisker (vibrissae) follicles, teeth, taste papillae, the intestine lumen, or salivary glands. In the developing vertebrate head, these tissue cross-talks employ neural crest-derived mesenchyme with the epithelium, myogenic progenitors, or cartilage. Critical steps in whisker follicle (WF) development are very similar to normal hair follicle (HF) development ^1^, which has been studied much more extensively. HF morphogenesis is initiated by thickening of the epithelium forming a placode (Pc), which subsequently induces condensation of underlying fibroblasts termed dermal condensation (DC) ^2^. Beta-catenin-dependent WNT signaling is essential for Pc induction ^3^, which initiates the EDA pathway ^4^, FGF20 ^5^, and SHH signaling ^6^ in the epithelium. Abrogation of any of these signaling pathways leads to the arrest of HF development. However, the succession of initial inductive steps in HF formation is still unclear. Pc formation requires hitherto unknown primary signals from dermal fibroblasts (Fb) at the pre-DC stage that is characterized by expression of *Foxd1* ^7^ and *Prdm1* ^8^. Recent studies employing single-cell transcriptomics helped elucidate the molecular determination of Pc and DC induction. Mok et al. ^7^ described a wave of transcription factors that define cell fate trajectories, starting in dermal fibroblasts underlying the epithelium before Pc induction (pre-DC stage), towards DC and the late DC stage when Pc invagination commences. For instance, expression of *Foxd1* transcription factor was detected in pre-DC fibroblasts that are also characterized by expression of *Twist2, Lef1*, and *Smad3* ^7,9^. DC cells predominantly express *Sox2*, *Foxd1, Ptch1,* and *Bmp4* ^10,11^. On the other hand, DC initiation requires canonical WNT and FGF20 signaling in Pc, suggesting a reciprocal molecular crosstalk between the dermal mesenchyme and overlying epithelium ^7,10^. Transplantation experiments suggested that mesenchymal cells modulate WNT signaling in Pc by activation of RSPO1; however, RSPO1 is not sufficient for HF induction even in combination with BMP inhibition ^12^. Although these data comprehensively map molecular events during HF initiation, identification of the primary inductive signal for Pc induction coming from dermal fibroblasts is still missing.

Epithelial structures such as whiskers or teeth are formed at precisely defined locations. Whiskers are arranged in an accurate rectangular pattern in five rows, each containing four to nine follicles. Each whisker is innervated by a separate whisker-to-barrel circuit that is reflected in a clear topographic organization of the barrel cortex ^13^. Axons of the trigeminal nerve ending in the infraorbital nerve innervate each whisker germ in the maxillary prominence. It has been suggested that early innervation does not play a role in determining WF pattern as distribution of trigeminal axons is widespread and random in the maxillary process, and no one-to-one spatial correlation between nerves and WFs was found ^1,14^ . This view has been challenged by the observation of a regular pattern of the nerve plexus in the snout half a day prior to the pattern of WFs ^15,16^. However, these studies do not imply whether WFs develop normally *in vivo* in the complete absence of innervation. Although normal vessel ring organization in the whisker follicles of mice lacking trigeminal sensory innervation has been reported ^17^, a comprehensive description of the WF development was not studied in this context. Therefore, while existing literature points to a scenario with an unlikely role of innervation for the WF formation and patterning ^18–20^, a more direct and conclusive *in vivo* study is needed to resolve this issue.

Similarly, the influence of the trigeminal nerves for tooth development has not been conclusively demonstrated. Organ culture methods showed no involvement of trigeminal (TG) sensory innervation ^21^, and tooth germs can trigger their own innervation ^22^. Along these lines, diastema tooth primordia develop and subsequently disappear with no TG innervation ^23^. However, the existence of earlier pioneering axons in the TG system has been reported; therefore, tooth development might be initiated by interaction with the early arriving pioneering axons ^24^. Moreover, roles of neurogenic factors have been shown during tooth development ^25–27^. A surgical removal of (TG) axons innervating teeth led to tooth degeneration ^28,29^. Hence, cranial nerves might deliver morphogenetic signals that orchestrate tooth maintenance. The role of innervation during tooth development was also supported by Kaukua et al. ^30^, who showed that dental mesenchymal stem cells are recruited from the nerve-associated glial cell. Taken together, it has been suggested that innervation play a role in tooth development ^31^. However, in a model system where innervation is completely missing, tooth development has not been analyzed.

Thus, peripheral nerves and accompanying glia contribute to the mesenchymal tooth germ niche. Nonetheless, it is still unclear what cellular mechanisms regulate precise positions of whiskers in the snout. Turing reaction-diffusion mathematical model can be applied for oscillating tissue patterning that is orchestrated by diffusion of short-range activating morphogen that triggers expression of long-range inhibitor. Reaction-diffusion-based instruction of the epithelium by combinatorial WNT, FGF, and BMP signaling is capable of HF placode induction in a highly restricted regular pattern ^32^. However, a highly specific rectangular arrangement and relatively long distance among WFs within the developing snout may indicate that other molecular determinants play a role in spatiotemporal control of WF induction. This revives the previous speculations that an axonal network may be involved in WF positioning. In this report, we assessed the scenario that the neural network is implicated in the control of accurate geometrical arrangement of WFs.

Here, we generated *Meis2* conditional mutants using the Wnt1-Cre2 driver ^33^ targeting exclusively the neural crest-derived tissues, including the dermal mesenchyme and cranial nerves, without detectable recombination in the overlying epithelium ^34^. Although *Meis2* expression in the epithelium of Wnt1-Cre2; *Meis2* ^fl/fl^ (*Meis2* cKO) mutants was preserved, whisker development was arrested at the Pc induction stage. This is documented by failure in induction of expression of placodal genes such as *Shh*, *Edar* and *Lef1*. We further show that severely affected branching of trigeminal nerves innervating WFs cannot cause their developmental arrest because WFs form normally even in a complete absence of trigeminal nerves. Expression of *Meis2* in dermal fibroblasts is therefore necessary for induction of whisker placodes followed by dermal condensation. Protein expression analysis suggests that MEIS2 transcription factor is essential for triggering a primary inductive signal from the mesenchyme to the epithelium to initiate WF placode formation. While single-cell transcriptomics and immunohistochemical validations showed that *Meis2* operates upstream of *Foxd1*, primary pre-DC marker, we report a dispensable role of *Foxd1* during WF development. Overall, our data not only describe an essential role of mesenchymal *Meis2* for WF development but also document normal WF formation even in the complete absence of innervating axons.

## Results

### *Meis2* expression in cranial neural crest-derived cells is required for correct trigeminal nerve projections and whisker development

To study whisker morphogenesis, we used Wnt1-Cre2; *Meis2*^fl/fl^ conditional mutants, in which *Meis2* is deleted in neural crest-derived cells (hereafter, *Meis2* cKO). Micro-computed tomography (micro-CT) of E15.5 embryos revealed severely compromised development of whiskers (Figure 1A). Missing whiskers were also observed whole-mount embryonic heads stained with placodal marker SOX9 antibody after light-sheet microscopy scanning at as early as E13.5 (Figure 1B). It is noteworthy that we observed in some cases a small number of SOX9+ WFs in *Meis2* cKO. They most likely represented normal follicles that ‘escaped’ from Wnt1-Cre2-mediated deletion of *Meis2*. The number of WF ‘escapers’ in *Meis2* cKO snouts varied among samples. Some mutant embryos completely missed WFs, but some showed a reduction in their numbers. Overall, quantification of the WF number documented a reduction to 5.7 ± 2.0% at E12.5 and to 17.1 ± 5.9% at E13.5 in the mutants (n=at least 3 embryos for E12.5, n= at least 8 embryos for E13.5, mean ± sem). Analysis of placode thickness/follicle length and DC area revealed that these escaper whiskers have normal morphology (Supp Figure 1A, B). To analyze expression of *Meis2* in the snout, we performed fluorescent immunohistochemistry. MEIS2 protein was detected in the epithelium, in the most superficial dermis including dermal condensates (DC). In *Meis2* cKO, mesenchymal expression of *Meis2* virtually disappeared (Figure 1C, asterisk, Supp Figure 1C) while its epithelial expression was maintained (Figure 1C, arrowheads, Supp Figure 1C), which corresponds to tissue-specific Cre recombination of Wnt1-Cre2 ^33,34^. It is interesting that MEIS2 signal was also remarkably reduced around sole WF ‘escapers’ (Figure 1C, arrow) but we cannot exclude that, due to incomplete Cre-mediated deletion, a minor expression of *Meis2* below the immunohistochemical detectability threshold is sufficient for WF induction. To verify the tissue-specificity in the snout region, we crossed Wnt1-Cre2 mice with mTmG mouse strain, where only cells derived from the neural crest express membrane-localized GFP. Whole-mount immunofluorescence imaging confirmed the Wnt1-Cre2 activity in neural crest derivatives in the craniofacial area, midbrain, and dorsal spinal cord (Figure 1D). In the snout, Cre activity was restricted to dermal mesenchyme without a detectable recombination in the overlying epithelium (Figure 1E). Moreover, branches of trigeminal sensory nerves were labelled with GFP (Figure 1E, arrows), consistent with the previously known origin of trigeminal neurons mainly from neural crest cells ^35^. Whiskers are innervated by the axons of the infraorbital nerve, a branch of the trigeminal nerve. To examine the axonal outgrowths associated with whisker germs, we co-labelled the hair follicle marker SOX9 with the sensory nerve marker TRKA in whole-mount heads at E13.5. Figure 1F and Supp Video 1 and Supp Video 2 show that missing whiskers in mice lacking *Meis2* in neural crest-derived cells were accompanied by severely affected trigeminal nerve growth and branching. Because defects in nerve branching were not limited to the infraorbital nerve and were also observed in other trigeminal branches, this phenotype is most likely not caused by missing WFs themselves. Moreover, the size of the trigeminal ganglion and the nerve exit from the ganglion appeared grossly normal (Supp Figure 1D). These observations suggest that defects in the trigeminal nerve morphology occur during branching and fasciculation of axonal projections through the craniofacial region. Overall, these results indicate an essential role of *Meis2* not only in the development of the whisker follicles but also for the branching of trigeminal nerves that innervate them.

**Figure 1.**
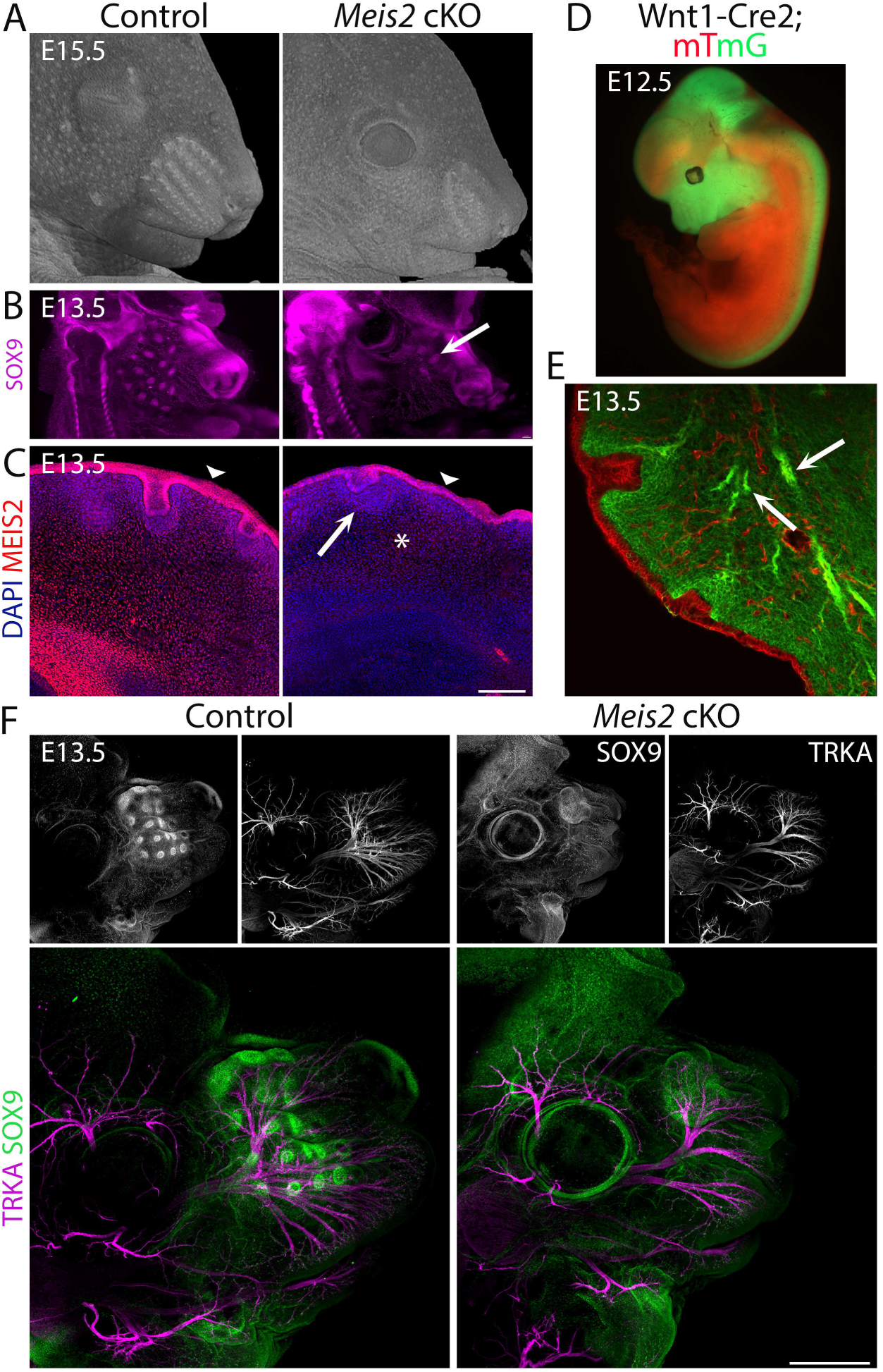
Severe whisker follicle development in *Meis2* cKO embryos. **(A)** Micro-CT images of control and *Meis2* cKO embryos at E15.5 showing aberrant whisker phenotype in *Meis2* cKO mice. **(B)** Light-sheet microscopy of Sox9 whole-mount immunostaining of WF. Arrow indicates example of an “escaper” whisker. **(C)** MEIS2 immunofluorescence on frontal frozen sections of E13.5 snouts showing MEIS2 expression in the dermis, DC, and epithelium including Pc. Arrowheads indicate epithelial expression of MEIS2 in both control and *Meis2* cKO mice. Arrow and asterisk show disappearing MEIS2 expression in the DC and dermis, respectively, in the mutant. Scale bar: 150 μm. **(D)** Wnt1-Cre; *mTmG* embryos showing Cre recombination specificity in the craniofacial area, midbrain, and dorsal spinal cord. Recombined cells are labeled by membrane-localized GFP in green, and non-recombined cells labeled by membrane-localized tdTomato in red. **(E)** Frontal sections of Wnt1-Cre; *mTmG* snouts documenting Cre recombination in the neural crest-derived mesenchyme, cranial nerve projections without recombination in the overlying epithelium. **(F)** Whole-mount staining of *Meis2* cKO heads at E13.5 with SOX9 and TRKA antibodies showing almost absence of WFs in and compromised branching of the trigeminal nerve in mutants. Scale bar: 500 μm.

Normal sizes of the escaped WFs in *Meis2* cKO mice at E12.5 and E13.5 (Supp Figure 1A, B) suggest that WF phenotype in the mutants is not due to a delay in their development since in this case WFs would appear smaller. In order to further exclude this possibility that WF formation is delayed in *Meis2* cKO mice, we analyzed whisker phenotype at later embryonic stages. Sections of E15.5 micro-CT images showed aberrant WF phenotype (Supp Figure 2A). Also at E18.5, the pattern of WFs were missing in the mutant (Supp Figure 2B). Staining of E18.5 snout section with placodal marker EDAR antibody revealed that, similarly to E13.5, a small number of escaped WFs in *Meis2* cKO with comparable depths to control WFs (Supp Figure 2C, arrows). However, this aberrant whisker phenotype persisted until E18.5. These data indicate that the whisker phenotype in the mutants is not due to a developmental delay but rather suggest a direct involvement of mesenchymal *Meis2* in mechanisms initiating WF formation. Interestingly, at E18.5 stage, hair follicles became detectable in the snout with a similar appearance in the mutants, suggesting a specific role of *Meis2* in WFs. This could be due to an early role of *Meis2* in the mesenchyme because HFs develop later.

### Trigeminal innervation is not required for WF development

Aberrant whisker development in *Meis2* cKO could be explained by an indispensable function of *Meis2* in mesenchymal cells or by insufficient trigeminal sensory innervation. The latter scenario takes into account the possibility that the nerve or nerve-associated Schwann cells produce factors that are necessary for WF formation. Thus, compromised branching of sensory axons in the *Meis2* cKO snout might result in insufficient secretion of crucial WF-inducing factors. In order to directly assess the function of sensory innervation in WF development, we analyzed *Neurog1*^-/-^ mice, which do not develop some cranial nerves, including trigeminal nerves, and therefore lack sensory innervation of the developing whiskers ^36^. Immunostaining of E13.5 100-µm thick snout cryosections for placodal marker SOX9, axonal marker TUJ1, and sensory nerve marker TRKA antibodies revealed that whiskers developed normally in *Neurog1*^-/-^ mutants even in the complete absence of innervating sensory axons (Figure 2A). Lack of any TUJ1-labelled axon fascicules in the proximity of the normally developed WFs in the *Neurog1*^-/-^ snout further excludes the possibility of non-sensory neurons triggering WF development in the absence of sensory innervation. In order to assess the role of sensory innervation for overall whisker patterning, we stained E12.5 and E13.5 snout whole mounts with SOX9 and EDAR antibodies, marking initial stages of WF development ^4^, i.e. placode formation. These results revealed essentially identical whisker patterning between control and *Neurog1*^-/-^ mice (Figure 2B). EDAR staining performed at E12.5 10-µm FFPE sections further confirmed normal placodal expression of EDAR, indicating normal initiation of the WF development (Figure 2C). WF formation was also validated by micro-CT at E17.5, showing normal progression of WF development at later stages (Figure 2D, top) even when trigeminal ganglia were absent (Figure 2D, bottom, arrows). Overall, these data unambiguously show that whisker formation does not require trigeminal sensory innervation and exclude the possibility that impaired nerves or nerve-associated cells in *Meis2* cKO resulted in impaired WF induction. Therefore, it can be confidently concluded that the dermal mesenchyme expressing *Meis2* is evidently critical for WF development.

**Figure 2.**
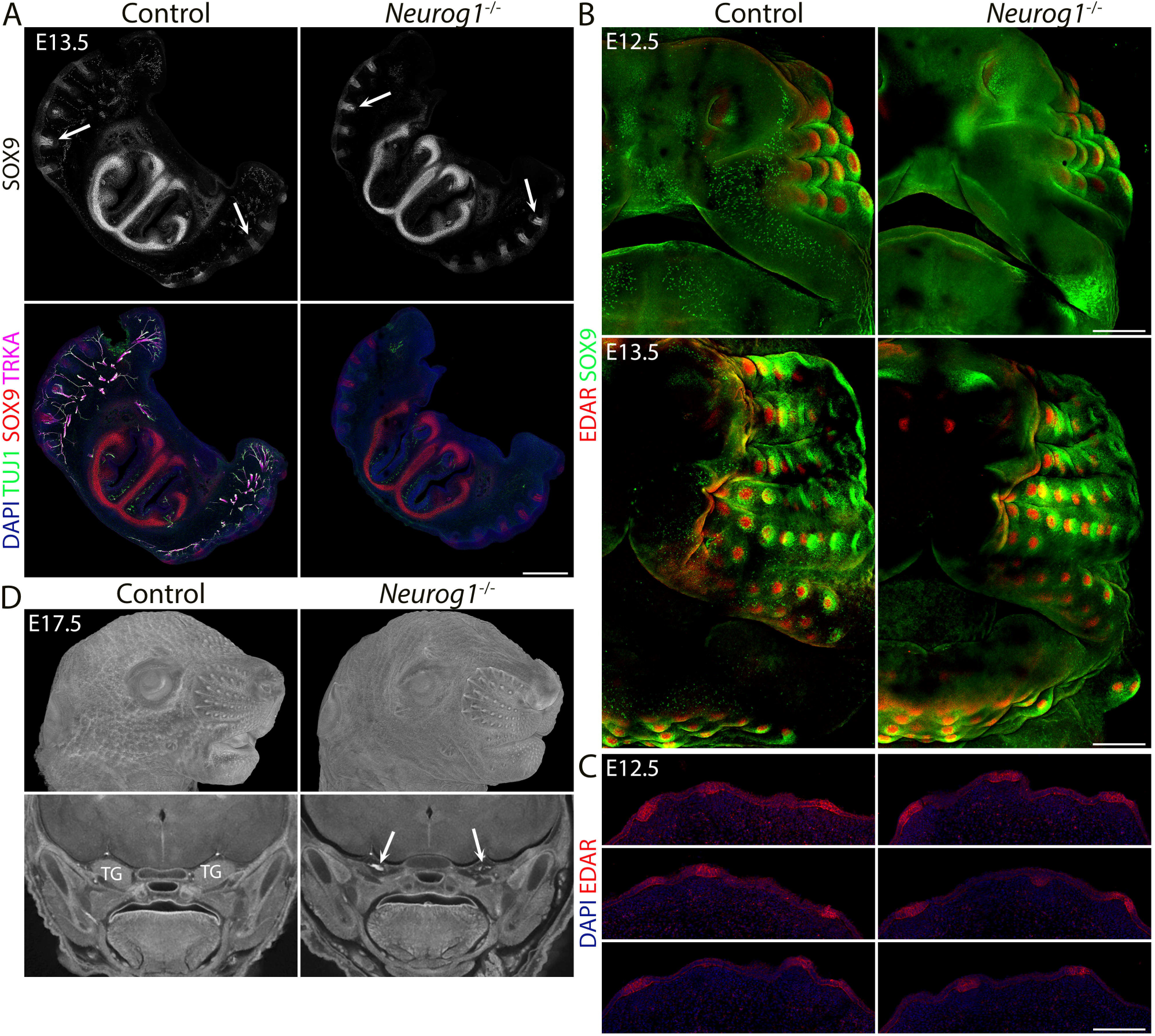
*Neurog1*^-/-^ (KO) embryos lack the trigeminal nerve, but WFs development is normal. **(A)** Triple immunostaining of 100-μm sections shows the absence of trigeminal nerve projections (TUJ1+, TRKA+) and normal WF (SOX9+) in *Neurog1*^-/-^ mice. Arrows in black and white images show examples of invagination of whisker placodes labeled by SOX9 antibody. Scale bar: 500 μm. **(B)** Whole-mount immunostaining of WFs with SOX9 and EDAR antibodies showing normal WF morphology and patterning in mutants at E12.5 (top) and 13.5 (bottom). Scale bars: 300 μm. **(C)** EDAR staining of 10-μm sections at E12.5 showing normal initiation of placode formation in *Neurog1* mutants. Scale bar: 150 μm. **(D)** micro-CT images reveal normally developed whiskers (top) at E17.5 in mutants while TGs are lacking (bottom, arrows).

In addition to whisker phenotype, we analyzed tooth morphology in *Neurog1*^-/-^ mice since the role of innervation for tooth development represents a long-standing controversy and has not been directly tested previously^21,23,24,31,37,38^. Interestingly, we did not detect any defects in tooth development (Supp Figure 3A), which was corroborated also by micro-CT images from E17.5 mice (Supp Figure 3B), reporting an innervation-independent formation of teeth in *Neurog1*^-/-^ mice.

### Mesenchymal *Meis2* operates upstream of epithelial EDAR signaling during initial steps of WF formation

Our data strongly suggest that *Meis2* expression in the mesenchyme is required for induction of placodal expression of *Sox9* in WF. Since a number of regulators of HF development were identified to function upstream of placodal *Sox9* expression, we tested these successive molecular events leading to WF induction. SHH release in basal placodal cells was shown to trigger *Sox9* expression in suprabasal placodal cells. This is achieved by asymmetric cell division resulting in suprabasal daughter cells with low WNT activity so that SHH can drive *Sox9* expression ^39^. We performed RNA in situ hybridization to evaluate *Shh* expression in *Meis2* cKO mice and control littermates at E13.5. These data revealed a remarkably decreased number of *Shh*-positive foci in *Meis2* cKO mice (Figure 3A, arrow), showing that placodal expression of *Shh* is regulated by mesenchymal MEIS2 protein. Again, we observed a small number of *Shh*-positive WF escapers (arrow). EDA/NFκB signaling in placodes has been shown to act upstream of Shh signaling before Pc invagination ^4^. We therefore stained whole mount E12.5 snouts from *Meis2* cKO mice and controls with SOX9 and EDAR antibodies. As seen in Figure 3B, EDAR protein was localized to epithelial WF placodes and overlapped with SOX9 staining in controls. In contrast, in *Meis2* cKO mutants, both EDAR and SOX9 were hardly detectable, documenting that *Meis2* mesenchymal function is upstream of EDA signaling in Pc. Whole-mount EDAR staining was confirmed on tissue sections which detected significantly reduced number of EDAR-positive foci in the *Meis2* cKO epithelium. Moreover, EDAR-negative regions within the mutant epithelium did not show any signs of morphological placode formation as inspected by DAPI staining (Figure 3C, Supp Figure 4A). Overall, these data demonstrate that mesenchymal *Meis2* operates at very early stages of WF formation in which it participates in forming a primary inductive mesenchymal signal for EDAR+ placode formation. This early arrest in mutants affects all subsequent steps of WF development including *Shh* and *Sox9* expression, and Pc and DC formation.

**Figure 3.**
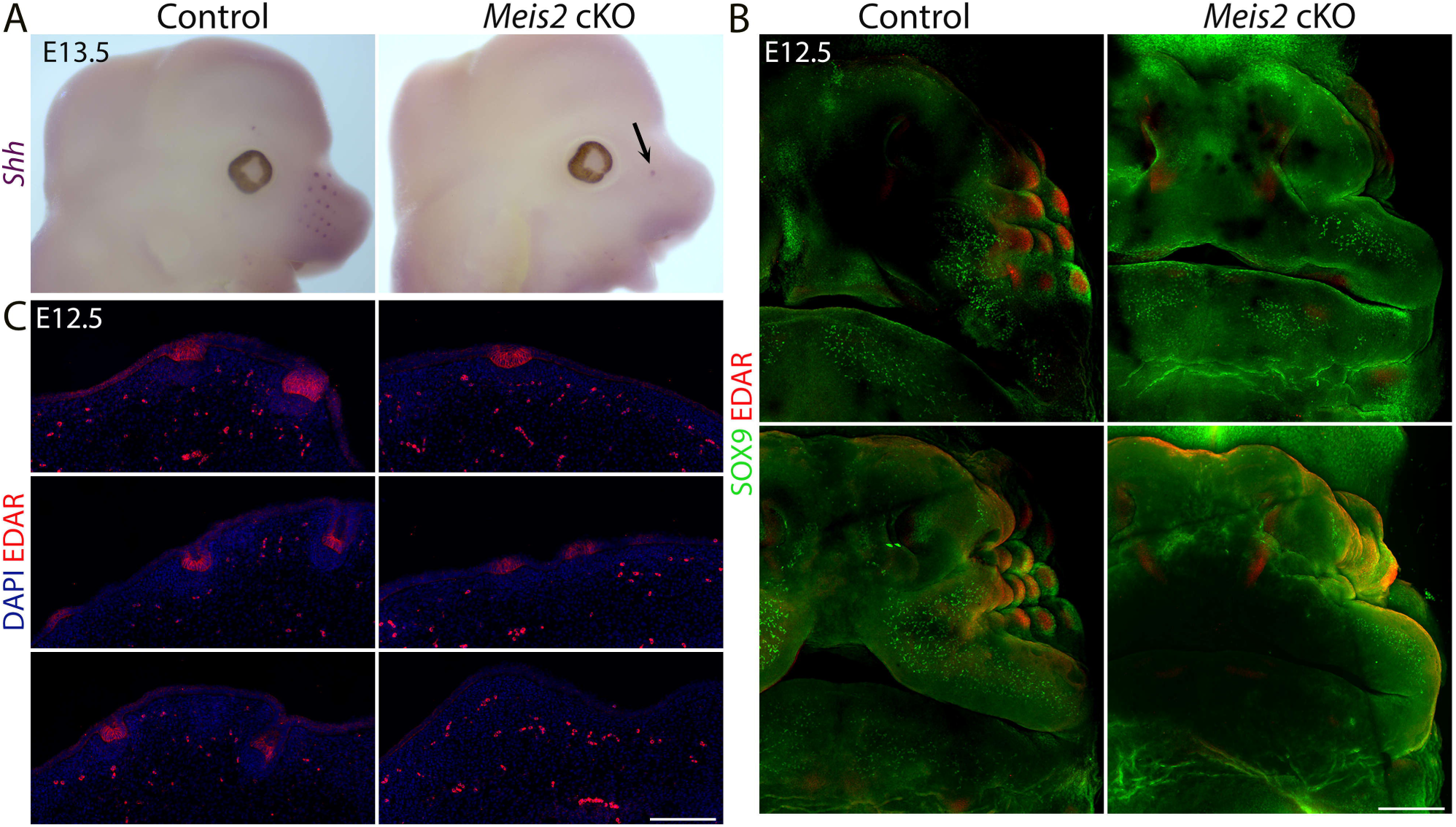
Induction and progression of WF development is compromised in *Meis2* cKO. **(A)** Whole-mount *in situ* hybridization of *Shh* mRNA documenting loss of WFs in mutants. Arrow shows an escaper whisker. **(B)** Whole-mount immunostaining of WFs with SOX9 and EDAR antibodies showing absence of WFs in *Meis2* cKO at E12.5. Two examples for each genotype are shown. Scale bar: 300 μm. **(C)** EDAR staining of 10-μm sections showing placode formation arrest in mutants. Scale bar: 150 μm.

### *Meis2* deletion in NCC-derived cells affects epithelial but not dermal WNT signaling

It has been shown that WNT/β-catenin signaling acts upstream of EDA/NFκB signaling as the earliest molecular pathway identified during HF formation ^4^. Epithelial WNT activity is required for activation of dermal canonical WNT signaling, while dermal WNT signaling reciprocally affects patterned WNT activity in the epithelium ^40^. We therefore assessed canonical WNT signaling in *Meis2* cKO snouts by several types of WNT signaling readouts. Firstly, we used LEF1 immunofluorescence because its expression has been widely used as proxy for WNT signaling in HF ^41,42^ and *Lef1* is also required for whisker formation ^43^. Matos et al. ^42^ showed that dermal cells expressed *Lef1* broadly whereas in the epithelium, *Lef1* expression was concentrated in Pc cells compared to inter follicular regions. During Pc downgrowth in HF, basal Pc cells displayed LEF1 signal similarly to Shh and EDAR, in contrast to suprabasal placodal cells lacking LEF1. This is consistent with the view that high WNT activity is seen in the basal cells whereas suprabasal cells are WNT low ^42^. In WF, we similarly detected robust LEF1 signal in Pc that was remarkably lower in interfollicular regions (Figure 4A, Supp Figure 3B, C). However, in distal snout regions with no WFs, the epithelium showed confluent LEF1 signal (Supp Figure 3C) suggesting that LEF1 signal is regionally dependent Pc indicator. In contrast to the epithelium, DC underneath invaginating placodes consistently displayed decreased LEF1 staining compared to peri-DC neighboring regions (Figure 4A, Supp Figure 4B). In *Meis2* cKO, we rarely detected a patterned placodal increase in LEF1 staining in the expected sites of WF positions (Figure 4A, Supp Figure 4C). Quantification of LEF1-stained E12.5 10μm FFPE sections revealed a significant decrease in LEF1+ placodal sites per section from 2.89 ± 0.39 in control to 1 ± 0.82 *Meis2* cKO (mean ± sem, p = 0.0002, two-tailed t-test, n = at least 3 mice, 18 sections). This shows that failure of WF initiation in *Meis2* cKO is accompanied by missing expression of *Lef1* in Pc, indicating that canonical WNT signaling is not induced in the Pc epithelium. However, average LEF1 intensity in the escaped placodes of the mutants did not significantly differ from that of control (95.08 ± 6.47 %, p = 0.51, two-tailed t-test). On the other hand, LEF1 staining in *Meis2* cKO dermal fibroblasts was confluent and without down-regulation in presumptive DC sites, which again confirms early developmental arrest in mutant WFs. Analysis of LEF1 staining intensity in both DC (93.60 ± 6.77 %, p = 0.305, two-tailed t-test, n = 3 mice and 43 DCs for control and 17 DCs for mutants) and non-DC upper dermis (89.91 ± 13.19 %, p = 0.117, two-tailed t-test, n = 3 mice and 37 regions for control and 33 regions for mutants) again did not reveal any significant difference between control and mutant snouts. These data indicate that *Meis2* in the mesenchyme does not affect dermal WNT signaling but is required for induction of epithelial WNT activity during placode formation.

**Figure 4.**
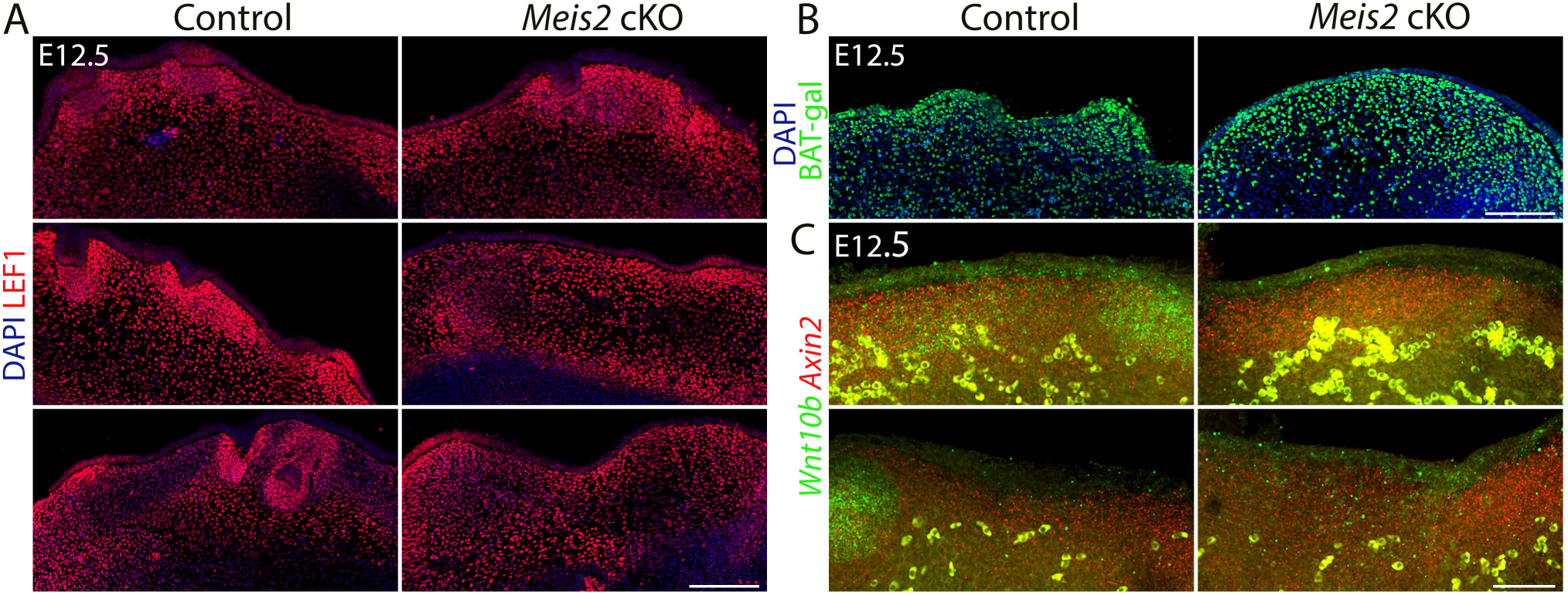
WNT signaling in the epithelium is affected by deletion of *Meis2* in the dermal mesenchyme. **(A)** LEF1 staining of 10-μm paraffin frontal sections of snouts showing abundance of LEF1 in the dermal mesenchyme, while in the epithelium, it is concentrated in placodes in regions of WF appearance. Similar to missing placodes, LEF1 is also lost in *Meis2* cKO epithelium. Scale bars: 150 μm. **(B)** beta-galactosidase immunostaining of sections from WNT reporter BAT-gal controls and *Meis2* cKO; BAT-gal mutants showing widespread galactosidase signal in the dermis and epithelium. Scale bars: 150 μm. **(C)** *In situ* hybridization HCR-FISH using probes for *Axin2* and *Wnt10b*. In the epithelium, *Wnt10b* is detected in placodes, and the signal is lost in *Meis2* cKO. Similarly, upregulation of *Axin2* is not observed in the expected WF loci. In the dermal mesenchyme, *Axin2* mRNA is detected throughout the dermis regardless of WFs. This pattern was not changed in *Meis2* cKO. Scale bar: 50 μm.

Secondly, in order to verify WNT activity dynamics in developing WF, we employed WNT reporter mouse strain BAT-gal ^44^ in which multiple TCF-binding sites promote expression of *beta-galactosidase* gene. We crossed this reporter mouse strain to *Meis2* cKO and stained snout frontal sections with an anti-galactosidase antibody. In contrast to LEF1 staining, BAT-gal reporter staining did not show consistent upregulation and concentration of WNT activity in Pc and no corresponding downregulation in DC in controls. It rather exhibited confluent expression in the dermal mesenchyme (Figure 4B) that was not changed in mutants. Thus, BAT-gal reporter therefore did not serve as a reliable WNT readout in developing WF in our hands, probably due to the high stability of galactosidase protein compared to RNA ^45^, which limited the analysis of the dynamic nature of WNT activity during WF development.

Thirdly, we tested the expression of selected WNT ligands in *Meis2* cKO snouts. *Wnt10b* expression is strongly activated in HF placodes ^46^ and it also is important for HF progression ^47^. Thus, *WNT10b* seems the most likely *Wnt* gene activating canonical pathway in HF which is routinely monitored by *Axin2* expression in many tissue contexts including HF ^10,39^. We performed HCR FISH to assess expression of *Wnt10b* and WNT target gene *Axin2* in snouts at E12.5. Similarly to LEF1 staining, we observed *Axin2* increase in wild-type Pc, while DC and dermal mesenchyme displayed uniform *Axin2* expression, which did not accord with decreased LEF1 in DC (Figure 4C, Supp Figure 4B, D). This might reflect differential responses of *Lef1* and *Axin2* expression to WNT activity in DC. In *Meis2* cKO, upregulation of *Axin2* in the epithelium was not detected which most likely reflected missing placodes. On the other hand, mutant dermal mesenchyme displayed broad *Axin2* expression which was similar to wild-type controls. *Wnt10b* transcripts were detected in normal Pc in controls while this expression disappeared in missing Pc in mutants (Figure 4C). Overall, these data suggest that mesenchymal *Meis2* does not affect WNT signaling in the dermis, whereas it is critical for the patterned WNT upregulation in the epithelium. Since WNT activation in the epithelium is one of the earliest steps of HF and WF development, this leads to loss of EDAR and *Shh* markers in *Meis2* cKO WF, and it confirms a very early role of mesenchymal *Meis2* during whisker development.

### *Meis2* regulates dermal cell proliferation and the transition of fibroblast cell fate to DC

We have recently reported single-cell RNA sequencing (scRNA-seq) datasets for mesenchymal cells derived from the cranial neural crest of E12.5 and E13.5 *Meis2* cKO embryos and their control littermates ^48^. Cell cluster analysis identified one cluster representing dermal fibroblasts. We subset and further re-clustered using the standard Seurat workflow (in Methods). We identified 6 clusters, where Cluster 2 was defined as DC by cell markers *Sox2*, *Sox18*, *Tbx18*, *Bmp4*, *Lef1*, and *Cdkn1a* (Figure 5A-B, Supp Figure 5A, Supplementary Data 1). The other clusters were determined to represent fibroblasts annotated by their markers in the heatmap and module scores generated for fibroblast (Fb), pre-DC, DC1, and DC2 markers (https://github.com/kaplanmm/whisker_scRNA/tree/main.) ^7,49^. (Supp Figure 5, Supplementary Data 1). Cluster 2 displayed low expression of Fb module score genes and high expression of DC2 module score genes, confirming again that cluster 2 represents developed DCs. Transcriptomic analysis showed that the cell number proportion in cluster 2 was substantially lower in *Meis2* cKO mice compared to control littermates (Figure 5C), reflecting the WF phenotype in the mutant. Interestingly, clusters 0 also contained a lower cell proportion in the *Meis2* cKO dataset. On the other hand, the cell number proportion in cluster 1 increased in *Meis2* cKO (Figure 5C). However, the cell identity of cluster 1 was unclear because it was marked by only 27 genes (Supplementary Data 1). Cluster 0 revealed GO terms related to cell cycle and chromosome segregation/organization according to GO analysis by clusterProfiler R package ^50^ using top 100 markers (Supplementary Data 2). Therefore, we conclude that cluster 0 very likely represents dividing cells. Lower cell number proportion in cluster 0 in *Meis2* cKO suggested a *Meis2*-dependent regulation of dermal cell proliferation. This hypothesis was tested by performing EdU chase experiments where EdU was injected at E12.5 and embryos were harvested after 2 or 18 hours. Indeed, the 2D area coverage of DAPI+ region by EdU+ region in the upper dermis of *Meis2* cKO snout decreased from 52.96 ± 2.05% to 42.38 ± 2.77% (mean ± sem, t-test p= 0.0057) in 2-hour pulse and from 52.17 ± 0.81% to 45.76 ± 1.85% (mean ± sem, t-test p= 0.0052) in 18-hour pulse (Figure 5D-E). These EdU experiments confirmed scRNA-seq data and indicated a *Meis2* function in proliferation of upper dermal cells which might contribute to the aberrant whisker phenotype of the mutant.

**Figure 5.**
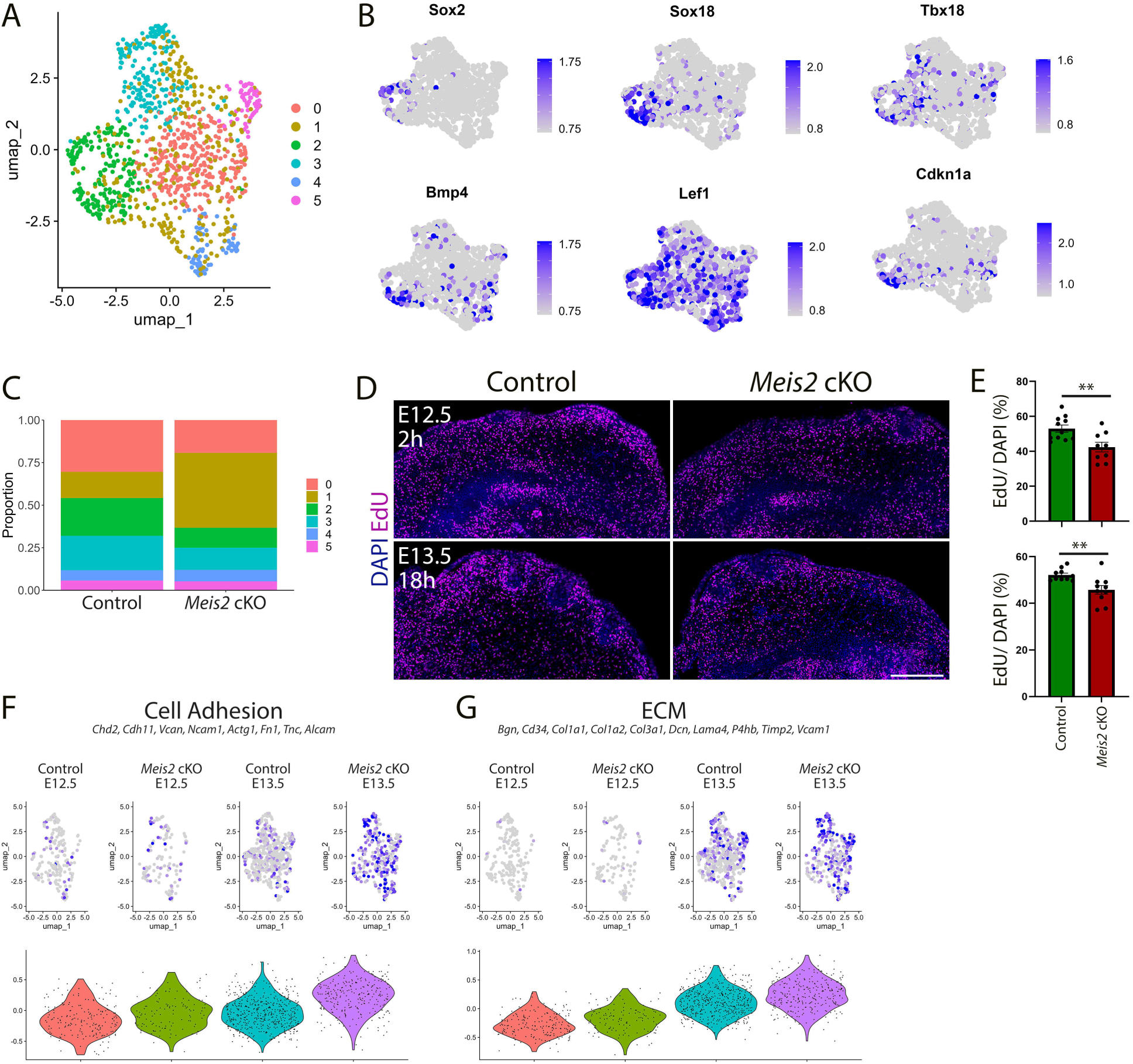
Dermal condensation and proliferation are reduced in *Meis2* cKO. **(A)** Uniform Manifold Approximation and Projection (UMAP) diagram of single-cell RNA-seq analysis of the dermal mesenchyme subset with 6-cluster resolution in which cluster 2 (green) represents dermal condensate (DC) of WFs. **(B)** UMAP representations of typical DC markers *Sox2*, *Sox18*, *Tbx18*, *Bmp4, Lef1*, and *Cdkn1* shown by FeaturePlots. **(C)** Relative cell numbers in 6 clusters showing a higher cell count in cluster 1 (nonspecialized cells) at the expense of clusters 0 (dividing cells) and 2 (DC). **(D)** Analysis of cell proliferation by EdU incorporation after a 2-hour pulse at E12.5 and an 18-hour pulse at E13.5. **(E)** Overall proliferation rate was reduced in *Meis2* cKO from 52.96 ± 2.05% to 42.38 ± 2.77% (mean ± sem, t-test p= 0.0057) in 2-hour pulse (top) and from 52.17 ± 0.81% to 45.76 ± 1.85% (mean ± sem, t-test p= 0.0052) in 18-hour pulse (bottom). Each data point indicates the average value for one section. n = 2 mice and at least 9 sections. **(F)** Increase of cell adhesion module score in mutants at E13.5 by scRNA-seq analysis. Module scores are generated by indicated genes. **(G)** Increase of extracellular matrix (ECM) module score in mutants at E13.5 by scRNA-seq analysis. Module scores are generated by indicated genes.

Because cell migration is a critical process for the migration of Fb cells into DC during cell fate acquisition ^51^, we plotted cell adhesion and ECM module scores, both of which were found to be increased in mutants (Figure 5F, G). Thus, such an increase in cell interactions might limit Fb migration efficiency by decreasing conductivity of the environment for cell migration and ultimately to decreased numbers of WFs in mutants. Similar results were found by Hudacova et al.^48^ in craniofacial mesenchymal cells in *Meis2* cKO mice. These increases in cell adhesion and ECM gene expression might also contribute to decreased TG axonal growth and branching described above (Figure 1, Supp Video 1 and Supp Video 2).

### *Meis2* controls expression of *Foxd1* in the dermis

Next, we performed differential gene expression analysis between the DC cluster of *Meis2* cKO and controls at E12.5 and E13.5. In this scRNA-seq dataset, *Foxd1* was found within the top genes downregulated in *Meis2* cKO mice (avg_log2FC = 1.0585, p_val = 4.55E-06) (Figure 6A, Supplementary Data 3). Interestingly, *Foxd1* expression was described as a hallmark of the earliest molecular changes in dermal fibroblasts prior to DC formation and defined the pre-DC stage in HF ^7^. Further, *Sox2*, a well-known DC marker, was also strongly downregulated in *Meis2* cKO (Figure 6A). To validate transcriptomic data, we analyzed FOXD1 protein expression in tissue sections of wild-type snouts at E12.5 and E13.5. Similarly to HF, *Foxd1* expression preceded *Sox2* also in WF which occurred before the Pc downgrowth into the dermal niche at E12.5. During initial Pc invagination and early DC stage at E13.5, *Foxd1* and *Sox2* expression overlapped (Figure 6B, arrows). In more progressed WF germs with deeply invaginated Pc, *Foxd1* expression remained in peri-DC regions rather than in DC that was marked by a strong SOX2 signal (Figure 6B, right panels, arrowhead). Next, whole-mount staining of FOXD1 in the whole snout highlighted dynamic WF development along the proximo-distal axis. It started in proximal columns at E12.5, in which WF formation was more progressed, while distal snout regions initiate WF slightly later (Figure 6C). By E13.5, 3-4 proximal columns displayed discernible WFs (Figure 6D). Strikingly, *Meis2* cKO snouts displayed no FOXD1+ WFs at E12.5 (Figure 6C). At E13.5, we detected around 4 proximal WFs in mutants while controls contained around 17 WFs on each snout side (Figure 6D). FOXD1+ WFs in *Meis2* cKO most likely represented ‘escaper’ WFs that were described above. Whole-mount FOXD1 immunostaining was further confirmed on tissue sections (Figure 6C-D, lower panels). Normal WF at E13.5 expressed *Foxd1* in DC or peri-DC and *Sox9* in Pc. In contrast, this progression of WF formation was halted in *Meis2* cKO (Figure 6C, D). These results suggest that MEIS2 is the key component of transcription controlling machinery in the dermal mesenchymal that initiates WF.

**Figure 6.**
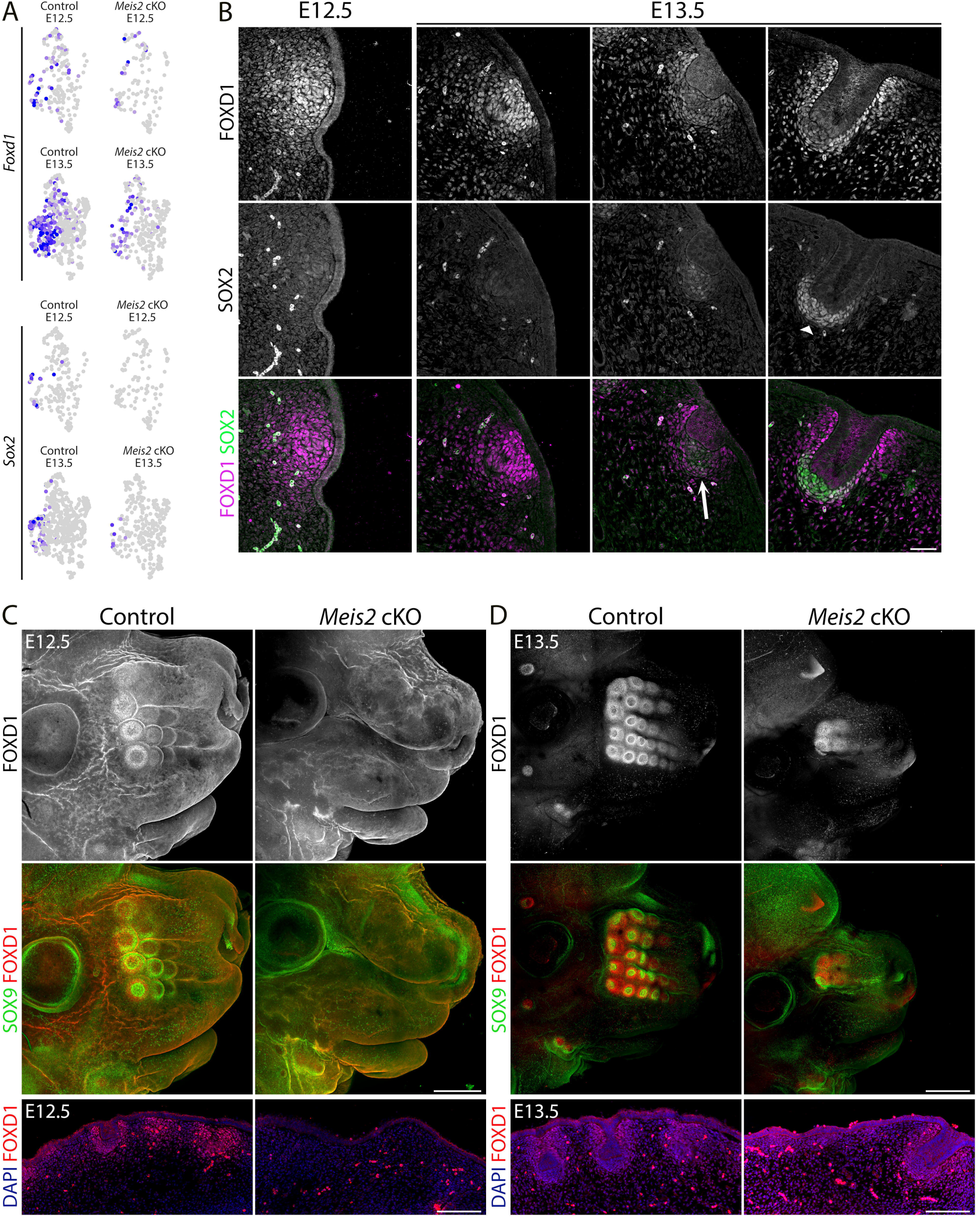
Expression of the earliest pre-DC and DC marker *Foxd1* is lost in *Meis2* cKO. **(A)** UMAP representations of decline in *Foxd1* (top) and *Sox2* (bottom) expression in DC cluster (#2) shown by FeaturePlots. Expression of both genes is significantly reduced in mutants. **(B)** Immunofluorescence of FOXD1 and SOX2 at initial stages of WF showing FOXD1 in the pre-DC stage while the expression shifts from DC to peri-DC regions during WF progression. SOX2 appears as a typical DC marker with no earlier expression. Scale bar: 50 μm. **(C-D)** Whole-mount immunostaining of FOXD1 and Sox9 of heads from controls and *Meis2* cKO at E12.5 **(C)** and E13.5 **(D).** It shows the absence of WFs including the pre-DC marker FOXD1 at the E12.5 stage when proximal columns of WFs have already formed. A few WF escapers develop in mutants at E13.5 in proximal regions while at least 6 WF columns have formed in controls. Scale bars: 400 μm (C) and 500 μm (D). Lower panels: FOXD1 staining in FFPE sections. Scale bars: 150 μm.

### *Foxd1* is dispensable for WF development

Because *Foxd1* expression in DC is strictly regionally specific and its expression is lost in *Meis2* cKO, we hypothesized that *Foxd1* transcription factor function is essential for WF formation. We therefore analyzed whiskers in embryos where *Foxd1* expression is ablated. We crossed heterozygous *Foxd1*-GFP-CreERT2 mice, in which GFP-Cre-ERT2 cassette was inserted into the initiation codon of *Foxd1* gene. Homozygous embryos are therefore null mutants for *Foxd1* gene. WF development was assessed in whole-mount-stained embryonic heads at E13.5 using SOX9, FOXD1, and TUJ1 antibodies. Figure 7A shows that SOX9 was detected in invaginating placodes, FOXD1 in surrounding DC mesenchyme, and TUJ1 in axonal projections innervating each WF. In parallel, we stained EDAR in WF placodes (Figure 7B). Interestingly, the number and morphology of WF in controls and *Foxd1* mutants were essentially identical. Thus, FOXD1 protein is a very important marker of the early DC cell fate in WF but does not have functional relevance for whisker formation despite its dynamic and regionally specific expression pattern around the developing WFs.

**Figure 7.**
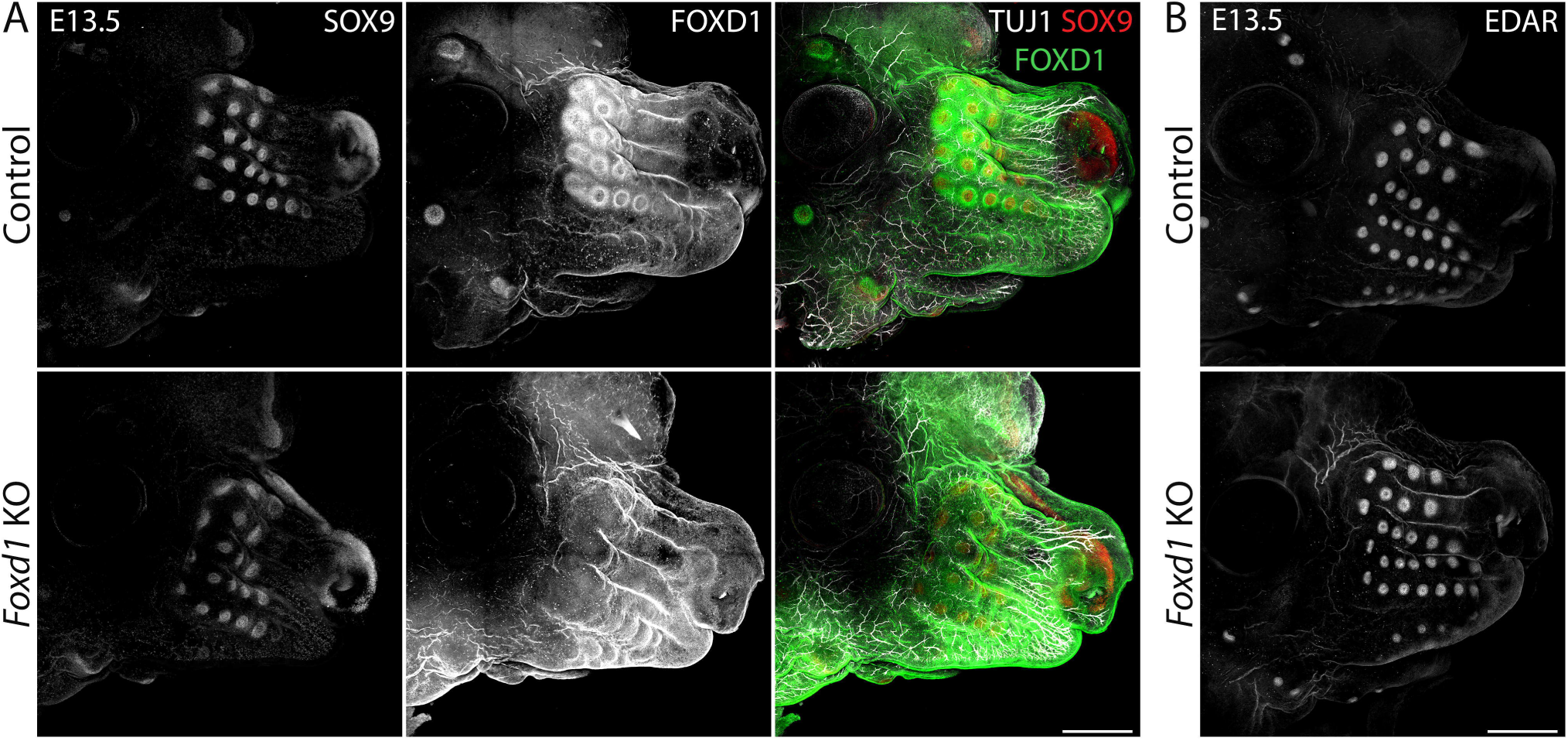
WFs normally develop in loss-of-function *Foxd1* mutants. **(A)** Whole-mount immunostaining of FOXD1, TUJ1, and SOX9 of heads from controls and *Foxd1*-null mutants at E13.5 showing normal formation of WFs in mutants in which FOXD1 signal disappears. Normal WF development is also reflected in normal WF innervation represented by TUJ1 staining. Scale bars: 500 μm. **(B)** Whole-mount immunostaining of EDAR confirmed normal Pc appearance in *Foxd1*-null mutants. Scale bars: 500 μm.

## Discussion

We have shown that mesenchymal expression of transcription factor *Meis2* is essential for the initial steps of mouse whisker development. Mouse embryos, in which *Meis2* was deleted in tissues derived from neural crest cells by Wnt1-Cre2 driver, lost the majority of whiskers. The aberrant whisker phenotype in the mutants is accompanied by markedly reduced trigeminal nerve branching in the snout. This suggests a novel function of *Meis2* in trigeminal axonal growth, branching, and guidance. It may also implicate that whisker formation, or WF positional determination, may depend on early sensory innervation. We report here, however, that whisker formation is entirely independent of sensory innervation. *Neurog1*^-/-^ mice completely lack trigeminal nerves, including their ganglions, but whisker development is not affected. Moreover, analysis of *Neurog1*^-/-^ mice at E17.5 reveals that sensory innervation is not required for the further growth and stabilization of WFs. The effect of trigeminal innervation deficiency on postnatal WF development could not be tested because *Neurog1*^-/-^ mice are lethal within a day after birth. All these experiments exclude the possibility that the whisker phenotype in *Meis2* cKO embryos is an indirect consequence of decreased trigeminal nerve branching. It rather indicates that mesenchymal *Meis2* is a pivotal regulator of WF formation.

Our observations suggest that nerve branching phenotype in the mutant is independent of trigeminal ganglia development or missing whiskers, as *Meis2* cKO mice showed normal trigeminal ganglia size and nerve exit from the ganglia as well as defective TG nerve branches that do not innervate whiskers. Thus, *Meis2* rather influences trigeminal branching and guidance through the snout. Description of the exact nature of factors controlling trigeminal nerve branching downstream of MEIS2 warrants further investigation. A number of molecular pathways, such as SLIT/ROBO, SEMA/NRP/PLXN ^52–56^, EPHRIN/EPH ^57^ signaling, have been implicated during TG nerve development, and *Meis2* might act upstream of mesenchymal expression of ligands and/or trigeminal expression of receptors of these pathways to control TG nerve development. One possible mechanism might involve SEMA3A, an inhibitory factor for trigeminal nerve growth during tooth innervation, and *Sema3a* null mice display hyper-branched trigeminal nerves ^55,58,59^. Our scRNA seq datasets indicated an increase of *Sema3a* expression in *Meis2* cKO mesenchymal cell populations (data not shown), which could explain decreased nerve branching in *Meis2* cKO mice and could be addressed in future studies.

Essential role of *Meis2* in the dermal mesenchyme during very early steps of WF development is supported by several key observations. Firstly, although epithelial expression of *Meis2* is preserved in *Meis2* cKO mice, thickening of the epithelium is rarely observed, suggesting that the release of WF placode-inducing dermal factors is under control of mesenchymal *Meis2*. It is widely accepted that HF or WF formation is initiated from the dermis by a ‘first dermal signal’ that has not been identified yet. Our data show that this process is dependent on *Meis2*. Secondly, the absence of *Meis2* results in the loss of early placodal markers LEF1 and EDAR and pre-DC marker FOXD1 at expected WF locations. Absence of patterned upregulation of LEF1 and EDAR was documented in HFs after blocking dermal WNT signaling ^40^. Interestingly, our analysis of WNT signaling readouts did now show remarkable changes in canonical WNT signaling in *Meis2* cKO dermis, indicating that normal dermal WNT signaling is not sufficient to trigger patterned LEF1 and EDAR upregulation in the epithelium when dermal MEIS2 is absent. These epithelial changes probably require additional *Meis2*-dependent intermediate steps, which might act either in parallel with, or downstream of dermal WNT signaling to induce placode formation. In the dermis, pre-DC marker FOXD1 expression was shown to be dependent on *Fgf20* expression in HF Pc and occurs after Pc formation ^7^. Our analysis of *Foxd1* expression in wild-type snout dermis revealed that *Foxd1* expression appears prior to morphologically distinguishable WF Pc. In *Meis2* cKO, FOXD1 signal in the dermis prior to WF Pc formation was not detected, which accords with single-cell RNA seq data. This all suggests that *Meis2* operates upstream of *Foxd1*. However, direct analysis of *Foxd1*-null mice did not reveal any phenotype in WF development. This could be explained by either no function of *Foxd1* in this process or by a possible functional compensation by its paralog *Foxd3*. Since *Foxd1* does not affect WF formation and therefore the absence of *Foxd1* cannot account for lacking WFs in *Meis2* cKO mice, loss of FOXD1 in these mice evidently is a result of lacking pre-DC and DC structures. Lastly, it has been shown in HF development that dermal fibroblasts proliferate extensively prior to converting to DC cell fate. DC cells, on the other hand, exit the cell cycle and become quiescent ^7,10^. We generally observed reduced proliferation in upper dermal cells in *Meis2* cKO mice. Lower cell proliferation could contribute to inefficient cell fate change of Fb to DC and might also lead to insufficient expression and/or release of Pc-inducing factor in the dermis of *Meis2* mutants. Overall, these observations strongly suggest that the absence of *Meis2* in mesenchymal cells affects earliest processes of WF development, even prior to Pc formation.

It is important to note that we observed small numbers (around 18% at E13.5) of WFs in *Meis2* cKO expressing normal placodal and DC markers. These rarely formed WF ‘escapers’ in *Meis2* cKO mice had normal WF morphology, because their placode thickness, follicle length, and DC size were comparable between controls and mutants at E12.5-E13.5. WF ‘escapers’ in mutants were probably not sensitive to Wnt1-Cre2-mediated deletion of *Meis2*. Although immunostaining of MEIS2 in snout sections indicated that *Meis2* expression around WF escapers was hardly detectable in mutants, a slight but undetectable expression of *Meis2* resulting from incomplete gene deletion might account for normal WF formation. On the other hand, it is also possible that *Meis1* paralogue can partially substitute for missing *Meis2* in its function in the dermal mesenchyme. Alternatively, *Meis2* function in WF development might be restricted to a short time window as each WF develops at different time points along the proximal-distal axis.

## Materials and Methods

### Mouse strains

Generation of the floxed allele of *Meis2* gene (*Meis2* ^fl/fl^) was described by Machon et al. (2015) ^60^ *Meis2*^fl/fl^ strain was crossed with reporter mouse lines R26R-*EYFP* ^fl/fl^ (RRID:IMSR JAX006148) before crossing with the Wnt1-Cre2 ^33^. This reporter enabled monitoring Cre recombination specificity. Wnt1–Cre2 (RRID:IMSR JAX:022137) was used for specific deletion of the *Meis2* ^fl/fl^ gene in neural crest cells. For lineage tracing experiments, we used Wnt1-Cre2 crossed with mTmG (RRID:IMSR JAX:007676). During embryo harvesting, Cre recombination was checked by monitoring EYFP fluorescence. In very rare cases (1-2 embryos in 10 litters), Cre activity in the whole body was observed, indicating germline recombination. Such animals were removed from the analysis, and their parents were removed from crossing schemes. Neurogenin1 knockout mice were purchased from Jax Lab (RRID:IMSR_JAX:017306). *Foxd1*-GFP-Cre-ERT2 mice were purchased from Jax Lab (RRID:IMSR_JAX:012464). The presence of the vaginal plug was regarded as the embryonic day 0.5 (E0.5). Pregnant mice were anesthetized using isoflurane gas inhalation and euthanized by cervical dislocation. Embryos were dissected and placed in ice-cold PBS. All mice were genotyped as previously described ^60^. *Neurog1* and *Foxd1* mutant mice were genotyped according to the Jax protocol. All procedures involving experimental animals were approved by the Institutional Committee for Animal Care and Use (permission #PP-10/2019) with every effort to minimize suffering and the number of animals used. This work did not include human subjects.

### Immunohistochemistry and image processing

All embryos used for staining were harvested at either E12.5 or E13.5 embryonic stages, and heads were fixed in 4% paraformaldehyde (PFA) overnight at 4^0^C. After PFA fixation, samples were prepared in three ways. (1) For FFPE staining, after dehydration in ethanol and xylene, samples were embedded in paraffin, and frontal 10-µm sections were prepared. Followed by stepwise rehydration, antigen retrieval in 0.1 M citrate buffer, pH 6.0, under pressure boiling for 12 min was carried out. (2) For cryosection staining, after dehydration in 30% saccharose for 24 hours, samples were embedded in OCT, and 100 µm cryo-sections were obtained. (3) For whole mount staining, whole heads were directly processed to permeabilization and blocking after PFA fixation.

Tissues were blocked in 5% bovine serum albumin (BSA) in PBS with 0.1% Triton X-100 for FFPE sections or with 0.5% Triton X-100 for thick cryosections or whole mounts for 1 hour and incubated in primary antibody solution (1% BSA in PBS and 0.1% Triton X-100 for 10-µm sections or 0.5% Triton X-100 for 100-µm sections and whole mounts). Primary antibody incubation was overnight for FFPE sections, two nights for cryosections, and three nights for whole mount staining at 4°C. Primary antibodies used: SOX9 (Merck Sigma, AB5535), TRKA (R&D Systems, AF1056), MEIS2 (GeneScript, custom), TUJ1 (R&D Systems, MAB1195), EDAR (R&D Systems, AF745), Lef1 (Cell Signaling, C12A5), β-galactosidase (Abcam, ab9361), SOX2 (R&D Systems, MAB2018), FOXD1 (Abcam, AB129324). Primary antibody solutions were washed out with (PBS, Triton X-100) and incubated in fluorescent secondary antibody solutions for one hour for FFPE sections, three hours for cryosections at room temperature or overnight for whole mounts at 4^0^C. Secondary antibodies: donkey anti-mouse, -rabbit, -goat with Alexa488, 568, or 647 fluorophores (ThermoFisher Scientific). Primary and secondary antibodies were used at 1:500 for thick section and whole mount staining and at 1:1000 for thin section staining. Immunofluorescent images were scanned by spinning disc confocal microscopy using an Olympus SpinSR10. Imaging was performed for whole mount stainings after tissue clearing for one hour at room temperature. Acquired z-stacks were maximum intensity projected using ImageJ. Figures were assembled using Photoshop.

For cell proliferation assays, EdU was injected intraperitoneally to pregnant females at E12.5 for 2 or 18 hours before harvesting (0.5 ml per mouse, 2.5 mg/ml). EdU Base Click kit was used for EdU detection according to manufacturer’s protocol on FFPE sections.

### Micro-CT

Embryos were fixed in 4% PFA for 2 days and soaked in Lugol’s iodine for several days. Scanning was performed on the instrument Bruker Skyscan 1272 with the resolution 3 μm.

### In situ hybridization

Antisense RNA probe for *Shh* was cloned into pGEM-T-easy vector (Promega) using primers: *Shh*-F TCACAAGTCCTCAGGTTCCG, *Shh*-R GGGCTTCAGCTGGACTTGAC. Antisense mRNA was transcribed with T7 polymerase. Whole-mount *in situ* hybridization was performed using standard protocols ^60^. Fluorescent in situ hybridization HCR RNA-FISH (Molecular Instruments) was performed according to manufacturer’s protocols.

### Single Cell RNA sequencing analysis

scRNA sequencing datasets are retrieved from integrated Seurat objects (GSE262468) ^48^ and processed with the standard Seurat workflow ^61^. Briefly, dermal fibroblast clusters were subset and split by condition, which was followed by SCTransform normalization, where mitochondrial, ribosomal, and cell cycle genes were regressed. Seurat objects were then integrated by using the IntegrateData function with 3000 integration features. For UMAP as a non-linear dimensional reduction embedding, 30 principal components were used. We used the Clustree package to determine the optimal resolution, which was set at 0.6 for the FindClusters function according to the developers’ instructions ^62^. SCT assay was used to generate feature plots and violin plots. Normalized and Scaled data of the RNA assay were used to extract cluster markers using the FindAllMarkers function and to generate heatmaps. The FindMarkers function of the Seurat package was used for differential gene expression analysis between conditions with default parameters. clusterProfiler (4.6.2) was used for GO analysis ^50^, where GO terms were extracted by the enrichGO function. This paper did not produce original codes. Codes used for analysis and lists of genes used to generate module scores can be accessed at https://github.com/kaplanmm/whisker_scRNA/tree/main.

### Quantification and Statistical Analysis

For placode thickness/WF length and DC size analysis, images of DAPI-stained 10 µm FFPE snout sections were captured with an Olympus SpinSR10 microscope and a 20X objective. With ImageJ software, maximum intensity projection was applied to z-stack images. DAPI staining was used to visually identify placodes and DCs. The distance from the outer layer of epithelium in the placodal region to the epithelial-mesenchymal boundary was measured. For 2D DC size, DCs were identified by densely accumulated cells underneath the placodes. The area of the region covered by these cells was measured using ImageJ.

The number of WFs was manually counted in embryos that were used throughout the study for whole-mount immunostainings (labeled by SOX9, EDAR, and/or FOXD1 antibodies) or *in situ* hybridization (*Shh* mRNA). The numbers of WFs at both sides of the snout were pooled. The decrease in the percentage of WFs in *Meis2* cKO mice was calculated in comparison to their control littermates. Values from each experiment were averaged to determine the overall decrease in the percentage of WFs.

For EdU/DAPI area measurements, images of EdU-chased and DAPI-stained 10 µm FFPE snout sections were captured with an Olympus SpinSR10 microscope equipped with a 20X objective. With ImageJ software, maximum intensity projection was applied to z-stack images. The dermal region between the epithelium and around a 100-µm distance into the dermis was used for analysis, while DC domains were excluded. The selected dermal regions were processed separately for thresholding steps to cover EdU+ and DAPI+ regions. Threshold values were determined visually to cover specific EdU and DAPI staining. The same threshold values were used for control and *Meis2* cKO mutants. Total area values for EdU and DAPI were used to calculate 2D EdU coverage of the DAPI regions (EdU/DAPI).

Statistical analyses were performed using GraphPad Prism 10 software. N numbers specified in the respective figure legends for each analysis. All the data are presented with mean ± sem (standard error of the mean). Two-tailed student‘s t-test was used to explore statistical differences between two experimental groups (Control vs. *Meis2* cKO).

## Acknowledgements

This work was supported by the Czech Science Foundation (grants 22-10660S, 23-06160S) and Masaryk University, Faculty of Medicine (MUNI/A/1598/2023). We thank the Microscopy Service Centre of IEM CAS and Light Microscopy Core Facility, IMG CAS, Prague, Czech Republic, supported by MEYS – LM2023050 and RVO – 68378050-KAV-NPUI. This work was also funded by MEYS Czech Republic (NanoEnviCZ, LM2018124) and EU Structural Funds Pro-NanoEnviCz (CZ.02.1.01/0.0/0.0/16_013/0001821).

## Author contributions

MMK designed and performed experiments, bioinformatics, and wrote the manuscript. EH performed experiments and bioinformatics. MM performed experiments. HT performed experiments. JK designed experiments. OM designed and performed experiments, wrote the manuscript, and provided funding. The authors declare no competing interests.

Supplementary Figure 1. **Normal sizes of escaper WFs and TGs in *Meis2* cKO mice. (A-B)** Quantification of 2D DC area (**A**) and placode (Pc) thickness of non-invaginated or length of the invaginated follicles (**B**) showing comparable values between control and *Meis2* cKO at E12.5 (left) and E13.5 (right). N = 4 embryos for E12.5 and 3 embryos for E13.5, and at least 30 WFs for controls and 10 for *Meis2* cKO mice. P-values for all graphs are larger than 0.05, student t-test. Data are presented as mean ± sem. **(C)** 3 examples of 100-μm frozen frontal sections through snout stained with MEIS2 antibody showing disappearing MEIS2 staining in the *Meis2* cKO dermis. Scale bar: 300 μm. **(D)** 100-μm frozen sections through TG ganglia stained with TUJ1 and TRKA antibodies showing non-defective TG ganglion size or exit of the TG nerve from the ganglia. Scale bar: 250 μm.

Supplementary Figure 2. **Whisker phenotype persists at E18.5 *Meis2* cKO mice. (A)** Micro-CT images through the snout of E15.5 control and *Meis2* cKO mice showing abnormal whisker phenotype in the mutant. Arrows show some escaper WFs. **(B)** Brightfield images of control and *Meis2* cKO at E18.5. **(C)** 2 examples of 100-μm frozen sections of snouts from E18.5 control and *Meis2* cKO stained with DAPI (blue) and EDAR (red) showing only a few whiskers in the mutant. In contrast, EDAR-labeled hair follicles appeared to be normal in the mutant. Arrows show some escaper WFs. Scale bar: 1 mm.

Supplementary Figure 3. **Normal tooth development in *Neurog1*^-/-^ mice. (A)** 100-μm frozen sections of molar teeth of control and *Neurog1*^-/-^ mice at E13.5 stained with SOX9, TUJ1, and TRKA antibodies showing the normal phenotype of the mutant. Scale bar: 150 μm. **(B)** Micro-CT images around the molar tooth level at E17.5 in *Neurog1*^-/-^ mice.

Supplementary Figure 4. **EDAR and LEF1 expression in *Meis2* cKO mice. (A)** 3 examples of 10-μm FPPE sections stained with EDAR antibody showing affected expression of EDAR in *Meis2* cKO mice. Arrows indicate EDAR-positive sites which are observed less frequently in the mutant snout. Scale bar: 500 μm. **(B)** 2 representative images of normally developed escaper WFs in the mutants by LEF1 staining. LEF1 expression is similar in just forming placodes (top) and invaginated (bottom). Additionally, LEF1 expression shows a decline in the DC regions compared to peri-DC. Scale bar: 30 μm. **(C)** 3 representative micrographs of LEF1-stained 10-μm FFPE snout sections showing normal expression of LEF1 in the dermis but the decreased number of LEF1+ placodes in the epithelium of the mutant snout. Scale bar: 500 μm. **(D)** *In situ* hybridization HCR-FISH using probes for *Axin2* and *Wnt10b* in control snout at E12.5 showing placodal expression of *Wnt10b* (green) and *Axin2* (arrows, red) as well as widespread expression of *Axin2* in the dermis. Scale bar: 60 μm.

Supplementary Figure 5. **Identification of DC cluster in scRNA sequencing datasets. (A)** Heatmap of top 10 markers for each identified cluster in scRNA-seq data. Each column represents individual cells, whereas each row represents indicated marker genes on the left. Genes in the black box are top markers of cluster 2 (DC). **(B)** FeaturePlots and VlnPlots of fibroblasts (Fb), pre-DC, DC1, and DC2 module scores in each cluster. See Methods for selected genes used for generating module scores.

