## Supplementary figures and images for "Mesenchymal Meis2 controls whisker development independently from trigeminal sensory innervation"

### Supplementary Figures

#### Supp Figure 1

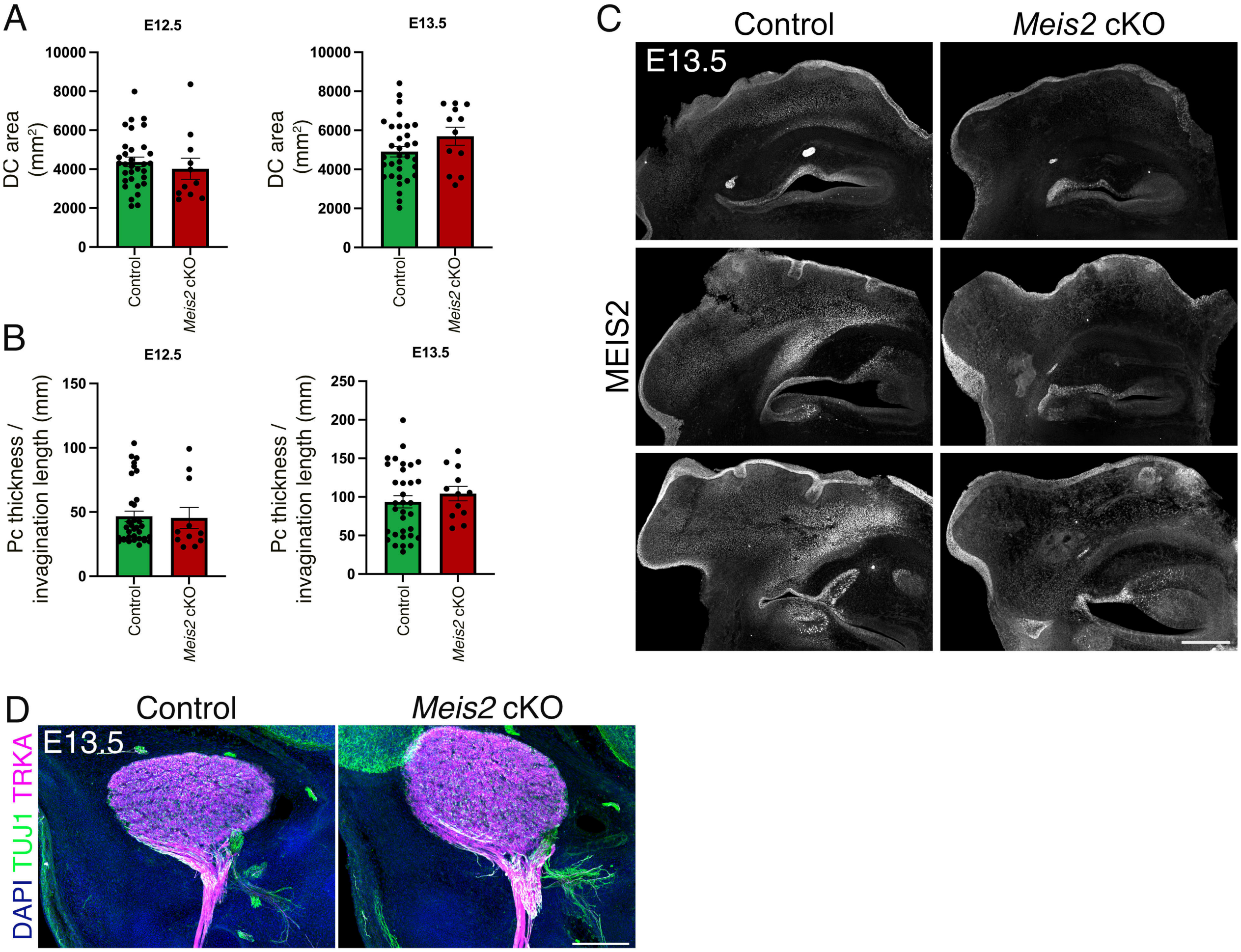

Supp Figure 2

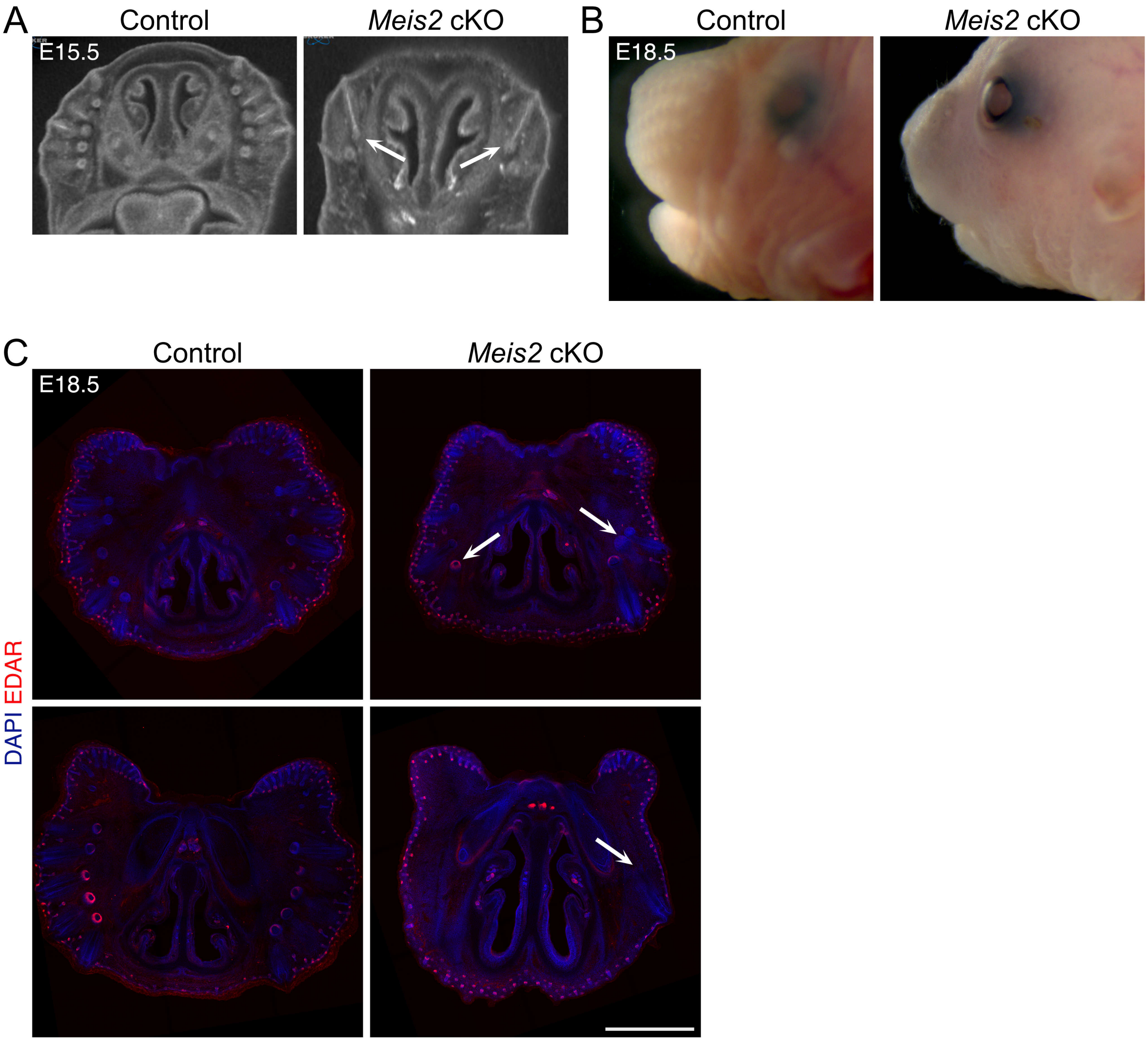

Supp Figure 3

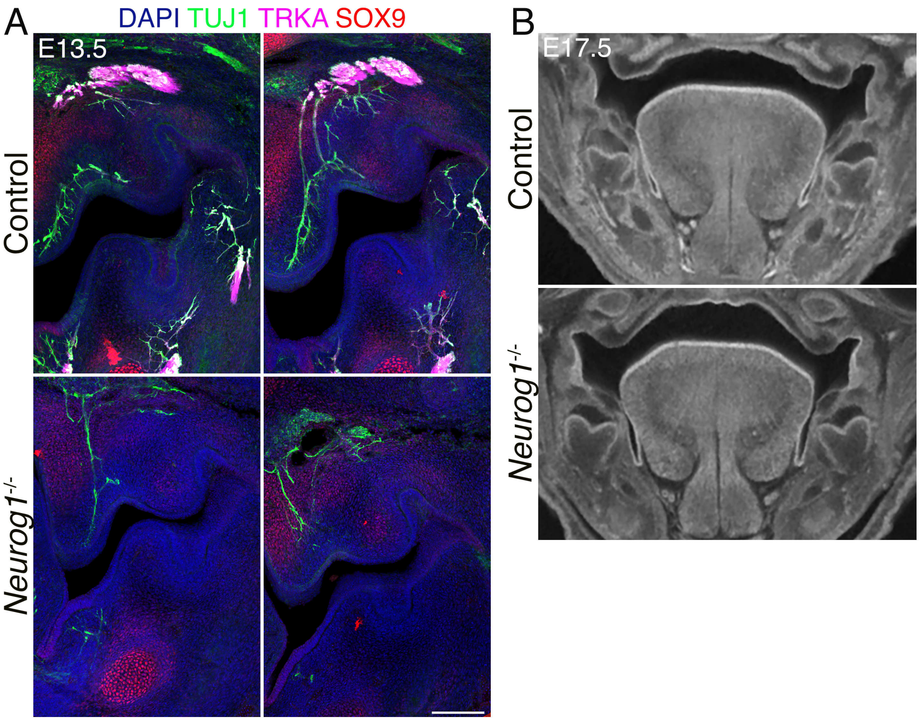

Supp Figure 4

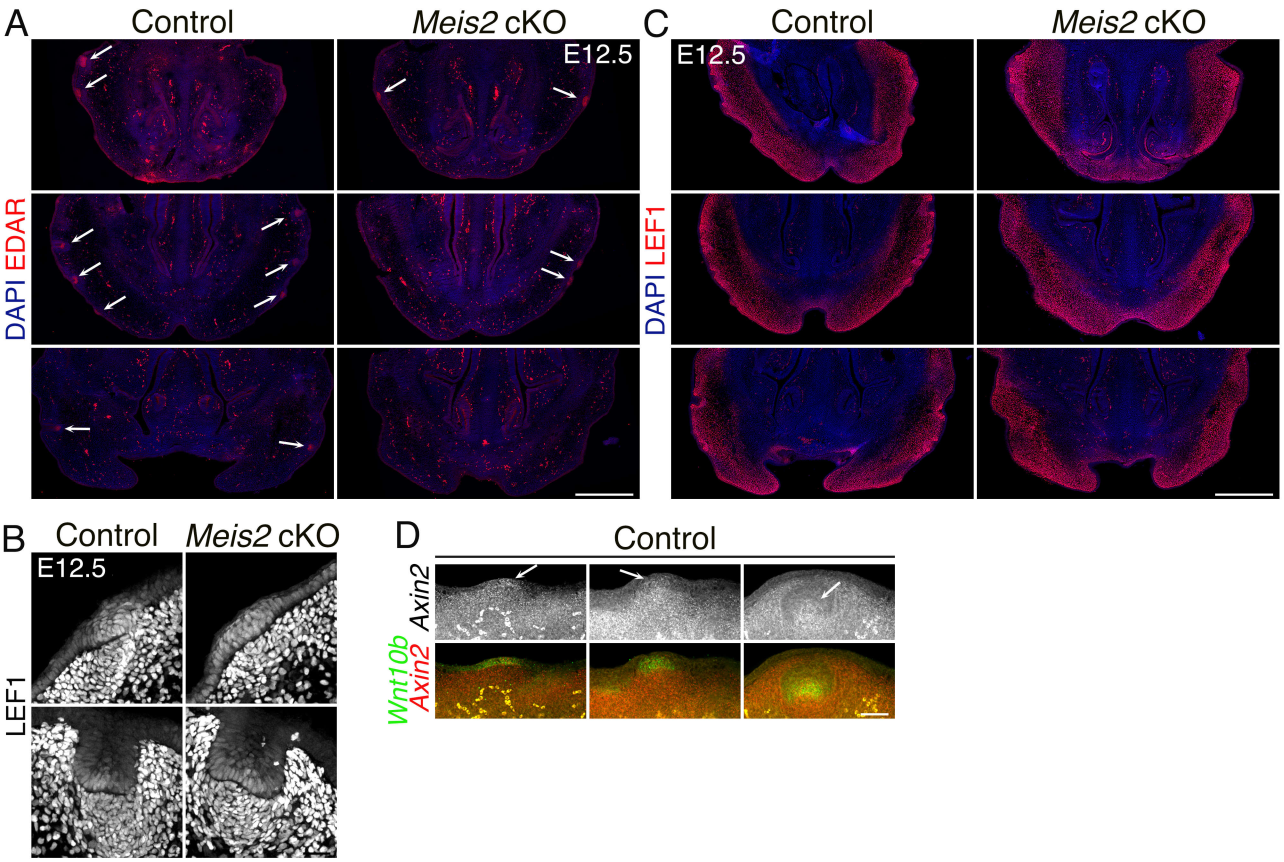

Supp Figure 5

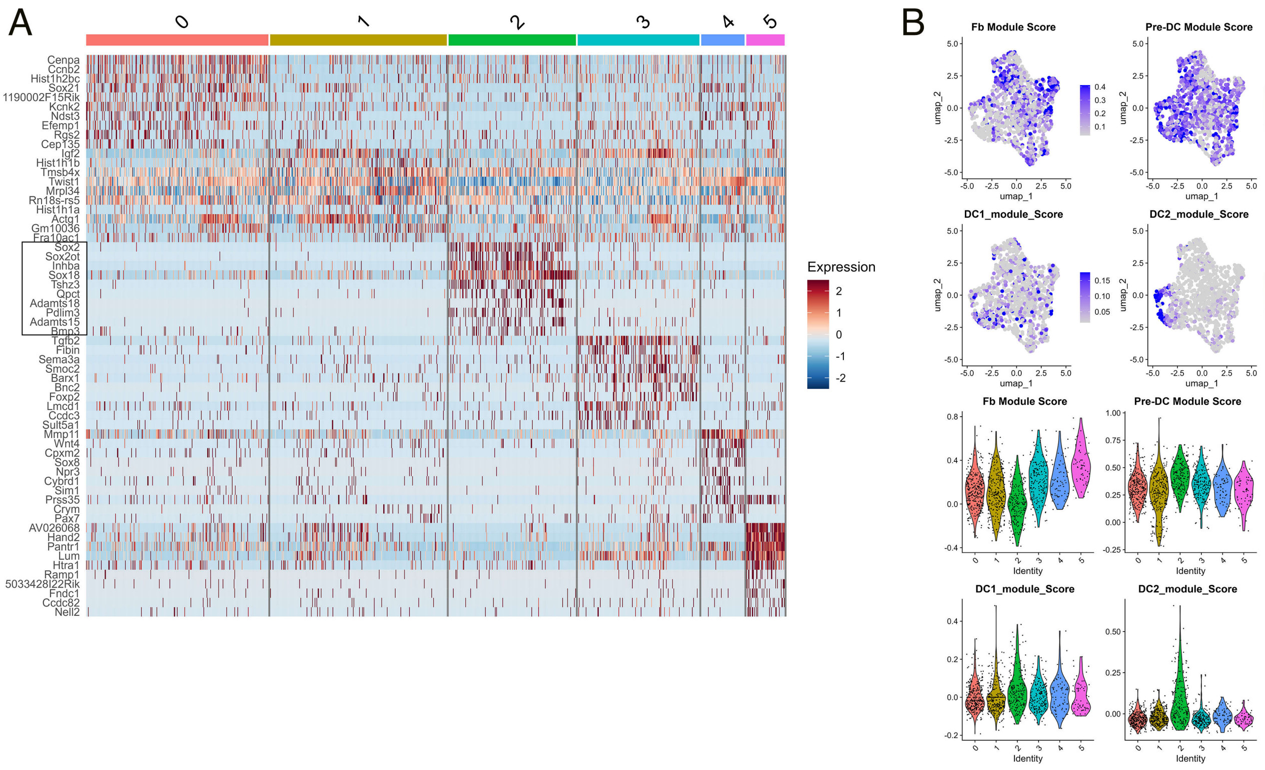
